# Assessing Drivers of Gender Balance and Racial Makeup of Editorial Board Members in Biomedical Engineering

**DOI:** 10.1101/2024.04.03.587972

**Authors:** Fariha Ahmad, Hannah Jackson, Matthew Kuhn, Katherine Danko, Jane Grande-Allen

## Abstract

Scientific journal editors serve as gatekeepers with the decision-making power of assigning reviewers to manuscripts. Serving as an editor is also an important stage in a young academic’s career progression, and an indicator of high regard/acceptance within one’s academic field. For both of these reasons, it is important to have representation with members of underrepresented groups serving in these roles. In this paper, we explore the gender and racial distribution among editorial boards for peer-reviewed scientific journals that are relevant to the field of biomedical engineering (BME). Further, we examine changes in these distributions from 2016 to 2021, amidst societal shifts catalyzed by movements such as #metoo and Black Lives Matter (BLM). Despite BME’s reputation for a relatively high percentage of female degree-earners, this study reveals stark disparities in gender and racial representation among editorial leadership positions. Through meticulous data collection and analysis of 75 BME journals— including 44 for which data from both 2016 and 2021 were analyzed—it was found that while the proportion of female editors increased over time, this proportion consistently fell short of the expected standard, which was based on current BME degree awardee values. Moreover, the percentage of Black editors remained stagnant. Correlation analyses between gender, race, and changes in journal impact factor (ΔJIF) revealed nuanced trends, in some cases showing that increasing ΔJIFs were associated with decreasing proportions of female editors. The study underscores the urgent need for changes in social and policy standards to address gender and racial inequities in BME [editorial] leadership, both of which will be necessary to foster greater diversity and inclusivity throughout the field.

## Introduction

The field of engineering revolves around finding creative solutions to problems. Since it has long since been recognized that having problem solvers from diverse backgrounds leads to higher quality solutions^1,2^, the field of biomedical engineering (BME) is often lauded given its relatively high percentage of female degree-earners^3^. Although the percentage of women earning degrees in BME is higher than in most other fields of engineering, metrics stretching beyond graduation statistics are often overlooked. For example, although 39 ± 6% of BME doctoral degrees are awarded by top-ranked institutions to women, only 17 ± 8% of faculty positions at those same institutions are held by women.^4^ Similarly, while people from underrepresented minority (URM) groups received 8 ± 8% of doctoral degrees awarded in BME, they made up little to none (4 ± 9%) of the faculty positions. Alarmingly, a trend of increased diversity in doctoral degree awardees but little change in faculty makeup is documented over decades in other science and engineering fields^4^, underscoring concerns of whether the diversity associated with the field of BME is overshadowed by positive trends in undergraduate degrees awarded.

One important effect of gender and racial distribution in BME can be ascertained by studying the distribution of both variables on editorial board makeup. Whether these positions are termed “editorial board members” or “associate editors,” people serving in this role—often heads of scientific labs or scientists with PhDs who are career editors—are responsible for evaluating the “fit” of submitted manuscripts for a journal, coordinating the peer review process, and making publication decisions^5^. However, when gender bias in editorial board makeup is not addressed by journals, resulting inequities in publishing can lead to barriers for women and URMs trying to reach higher levels of academic leadership.^6,7^

Although the need for hiring editors on the bases of gender distribution is recognized^8^, a study into whether or not these efforts have been made in BME-related journals has not been investigated. Additionally, society itself underwent rousing change in the latter half of the 2010s. Movements like #metoo and Black Lives Matter (BLM) swept across social media^9,10^, engendering further conversation revolving around the importance of diversity, equity, and inclusion (DEI) at all levels of society, including barriers of career progression in STEM.

The following study was completed to better understand gender and racial distribution in BME at the editorial board level. Additionally, in this study, we analyzed changes in editorial board makeup between 2016 and 2021, gauging variations that may have been made to editorial boards in the wake of the #metoo and BLM movements.

## Methods

### Data Collection

The first step of the data collection process was choosing the journal titles on which to perform gender and racial analysis. Multiple journal indices were considered, including Scimago journal rankings (SJR), Google Scholar, and Thomson-Reuter’s journal citation reports (JCR); JCR was chosen due to its impact factor data. This data was accessed through the Web of Science database provided by the Fondren Library at Rice University. Since the BME field is very interdisciplinary, we also considered a broad range of journals in which our Rice University bioengineering faculty have published. In 2016, journal titles in BME were initially selected based on their impact factor (impact factor > 1). In 2021, the list of journals was expanded to include additional high-impact journals that BMEs publish in, but are more general in scope. Journal titles that focus on publishing only review articles were omitted.

Journals store their editorial information on a variety of online formats; thus, the data was collected manually. Each journal title was looked up individually online, and the names of editors in the most universally standard positions (editor-in-chief, associate editors, editorial board members) was stored.

### Gender and Race Categorization

While gender is complicated and non-binary, the simplification that is assumed in this project is that gender is either female or male. With this simplification, the first gender identification method involved inference from first name and manual lookup. When the gender associated with a name could not be assumed with confidence, the editorial board member’s demographic information was searched for online and gender was confirmed by the pronouns used on their individual web pages. This method was tedious, and although pronoun information was available for almost all names, for some names this information could not be ascertained.

The second method of identification used social security data to predict the likelihood of a name being female or male. With the help of Dr. Fred Oswald (Department of Psychological Sciences, Rice University), RStudio (Posit Software, PBC, Boston, MA) was used to implement a code that would input name data and output a gender probability using the U.S. Social Security database. The code uses the proportion of female versus male to output one gender for each name. If the probability of “female” was higher than “male,” we assumed that the editor was female, and vice versa. In some cases, if the gender was undetermined using the R code, the editor was manually searched as described above. If the gender of the editor could not be determined using either method, the editor was excluded from later analyses.

A similar code was implemented to evaluate the probability of the racial background of the editors. Editors were divided into the following categories: White, Black, Asian, Hispanic, Native-American, or undetermined.

### Changes in Journal Impact Factor Analyses

To evaluate the relationship between changes in journal impact factor between 2016 and 2021 (ΔJIF), gender, and race, we first eliminated all editors who were listed multiple times in our data set. To do so, we ordered the ΔJIF from highest to lowest values to capture the largest ΔJIF, then removed any duplicated editors from the list such that only the most positive change in impact factor was recorded for a single editor. This process was repeated for both the 2016 and 2021 datasets. As with the previous analyses, if the gender or ΔJIF could not be assumed/calculated, the data point was disregarded.

### Statistics

Statistical analyses were completed using Microsoft Excel. Chi (χ)-squared analyses were used to determine significant differences between female and male ratios. The expected percentage of women was assumed to be 40%, based on recent data describing the proportion of women who receive doctoral degrees in biomedical engineering. All raw data were converted to percentages relative to the total number of editors. The null hypothesis was considered as rejected when p < 0.05, and the DOF = 1.

## Results

We analyzed a total of 44 journal titles in 2016 and 75 journal titles in 2021 (including the original 44 journals from 2016). In total, there were 1,608 editorships over multiple editorial positions with the majority being members of the editorial boards/associate editors.

### Trends in Gender and Racial Distribution

We first evaluated changes in female to male proportions across all journals over time. In 2016, women made up approximately 18.1% of all editorship positions. This percentage grew to 23.9% by 2021, but was still significantly less than was expected (p = 9.9×10^-4^).

When considering race, we first assessed broad changes in racial distributions over time.

We found no differences in racial proportions between 2016 and 2021, with approximately 57% of editor positions held by White individuals, 39% held by Asian individuals, 3% held by Hispanic individuals, and 1% held by Black individuals (Figure 1).

**Figure 1.**
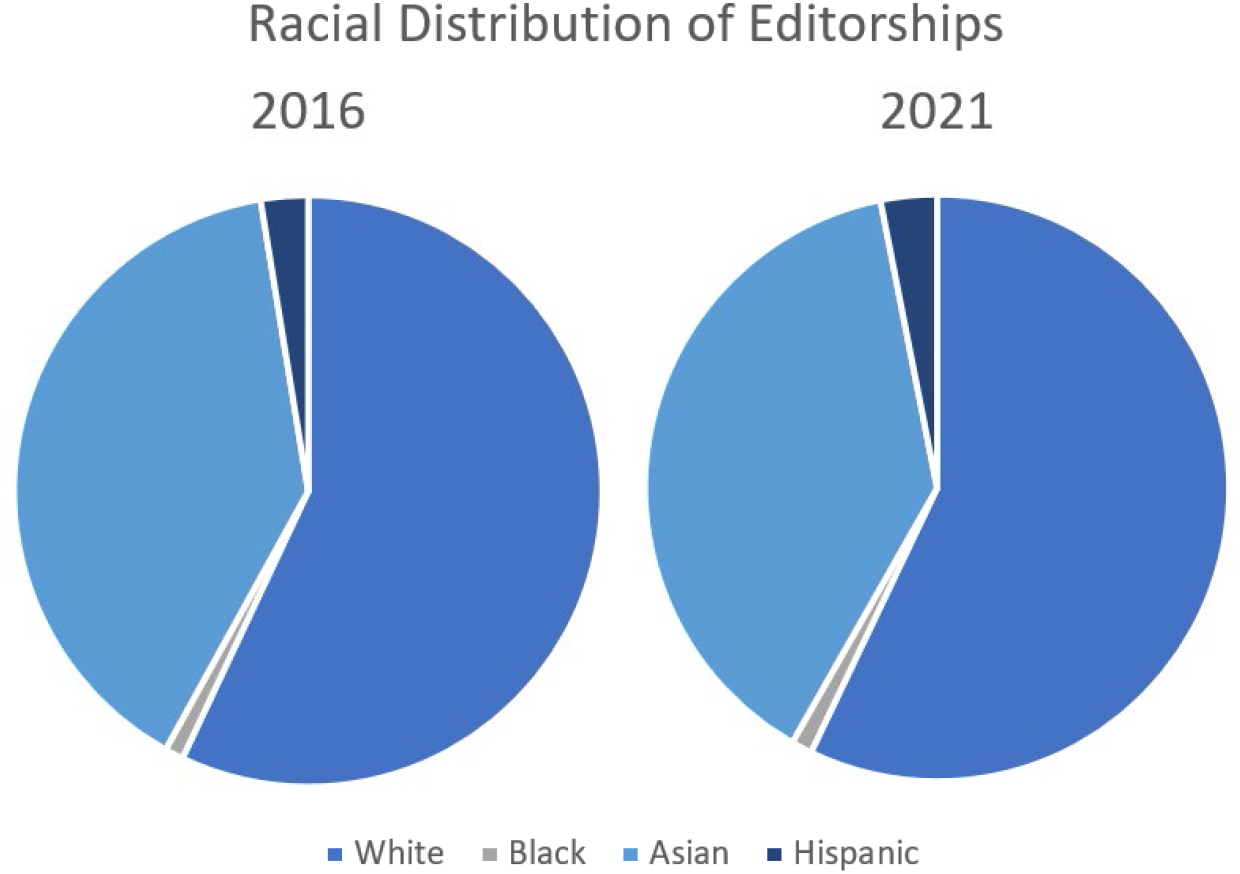
Racial Distribution of BME Editorships in 2016 and 2021

We then considered changes in proportions of women and men within racial subcategories over time. As shown in Table 1, while the percentages of women in each racial group increased over time, they never reached the expected percentage (40%).

**Table 1.** Gender Distribution in Racial Categories Between 2016 and 2021.

| 2016 |  | White | Asian | Black | Hispanic |
| --- | --- | --- | --- | --- | --- |
|  | % Men | 80 | 87 | 86.7 | 73.7 |
|  | % Women | 20 | 13 | 13.3 | 26.3 |
| | p-value | $4.5.9 \times 10^{-5}$ | $3.6 \times 10^{-8}$ | $5.2 \times 10^{-8}$ | $5.2 \times 10^{-3}$ |
| 2021 | % Men | 75.6 | 79.3 | 78.4 | 72 |
|  | % Women | 24.3 | 20.7 | 21.6 | 28 |
| | p-value | $1.4 \times 10^{-3}$ | $8.3 \times 10^{-5}$ | $1.8 \times 10^{-4}$ | 0.01 |

### Correlations Between Gender, Race, and Impact Factor

When assessing the ΔJIF, we found that overall, the percentage of women associated with journals that had an increase in journal factor between 2016 and 2021 was less than men. When comparing proportions of women and men, ΔJIF, and time, we found that although the percentage of women associated with journals that had positive ΔJIF increased between 2016 and 2021, the increase in proportion was limited by impact factor (Figure 2). The percentage of women associated with journals that saw an increase in impact factor increased until the ΔJIF reached approximately 4. For journals that had a ΔJIF greater than 4, the percentage of women associated decreased, including journals that saw a ΔJIF greater than 9.

**Figure 2.**
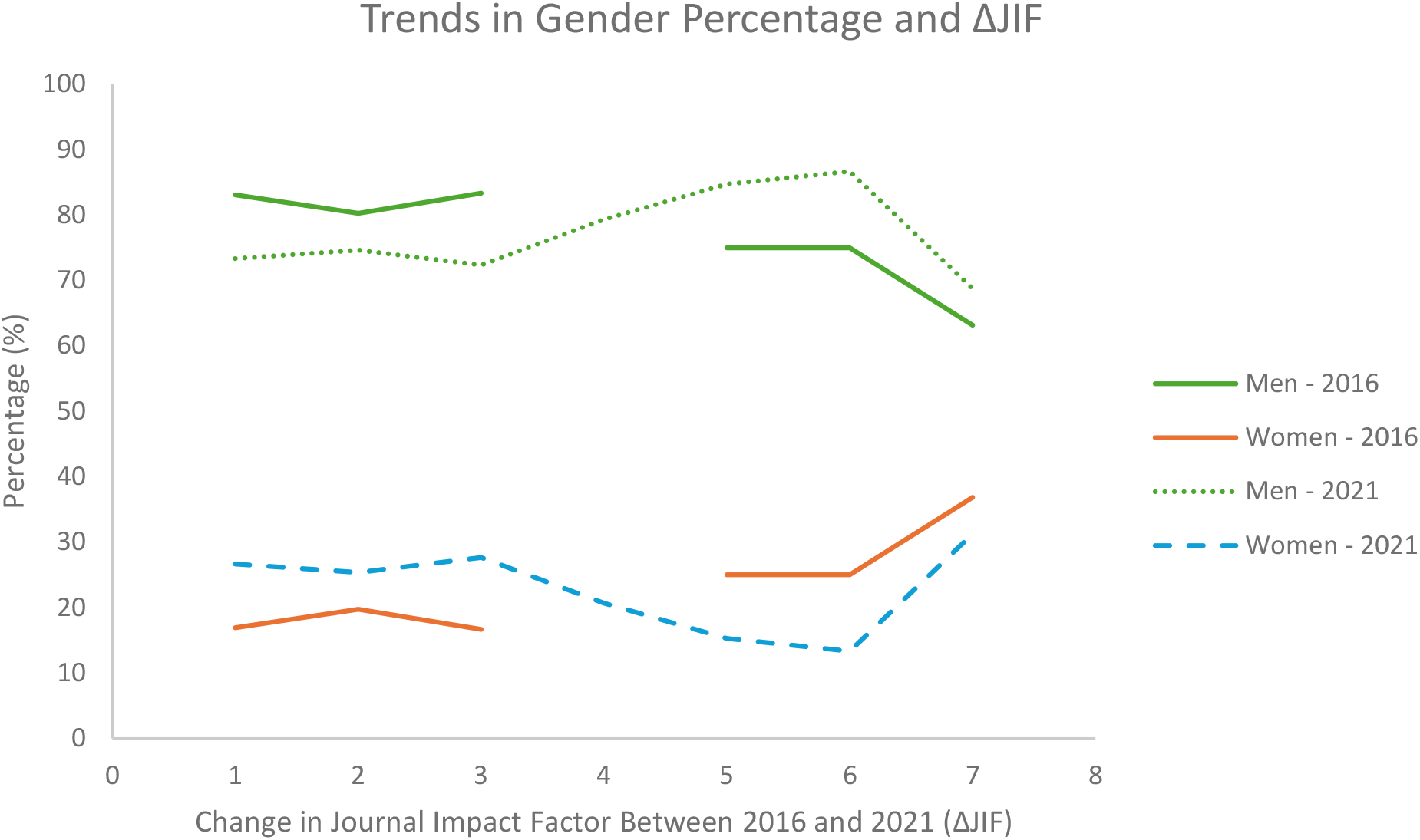
Trends in Gender Percentage and ΔJIF

We then considered the relationship between racial distribution and ΔJIF (Figure 3). We found that the percentage of White editors associated with journals that had a positive ΔJIF increased (from 60.5% to 62.4%) between 2016 and 2021, while the Hispanic editorship percentage decreased (from 3.1% to 1.8%). The percentage of Asian and Black editors associated with journals that had a positive ΔJIF stayed relatively similar (35% and 1%, respectively).

**Figure 3.**
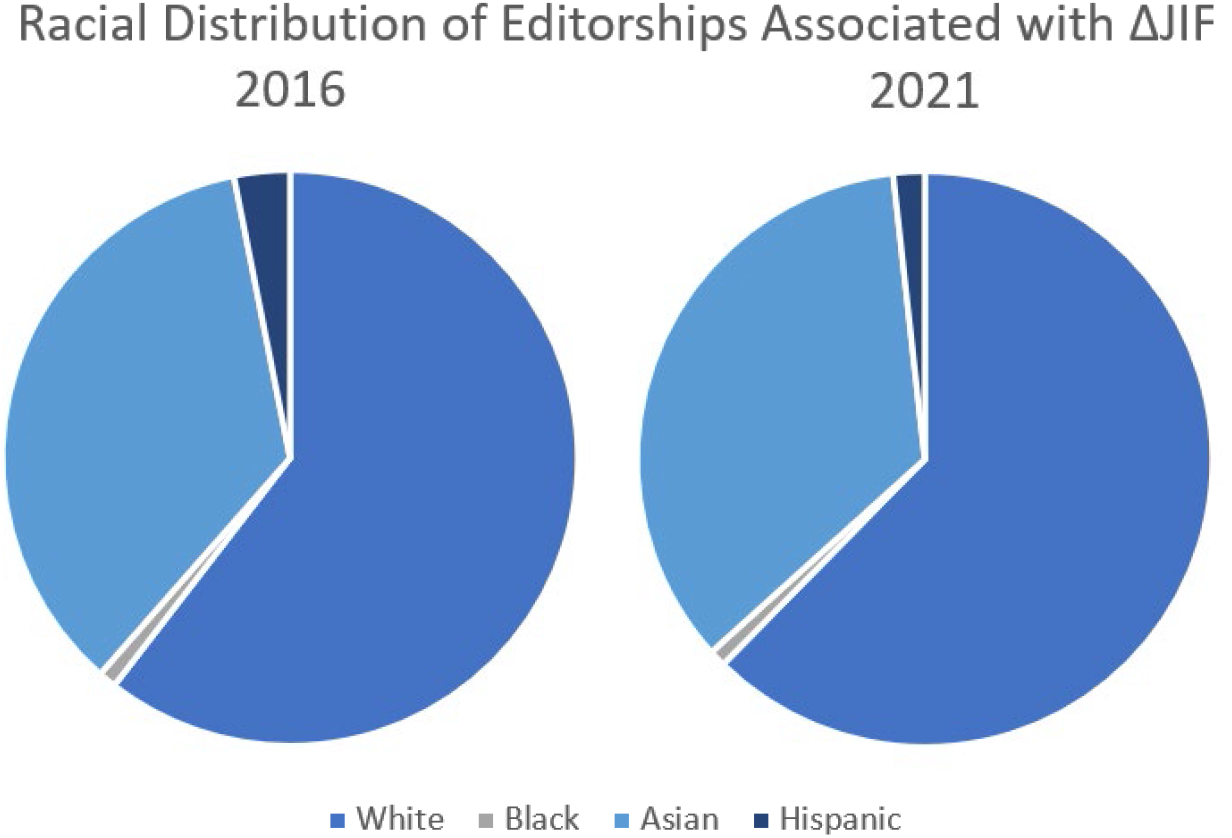
Racial Distribution of Editorships Associated with ΔJIF.

## Discussion

Journal editors play a significant role in the publishing process of manuscripts into scientifical journals; unfortunately, gender bias is also a major factor throughout this process. Correlations between the gender of editors and the gender of reviewers they assign^11^, editor gender and corresponding-author gender, and same-gender agreement between reviewer and editor have been documented^12^.

While BME is considered a field with higher diversity then other STEM and engineering fields, in this study we found that analyzing the gender and racial makeup of BME editorships quickly wore away the façade of BME as an equitable field, especially at higher leadership levels. Although the percentage of women holding editor positions at BME journals grew between 2016 and 2021, the proportion of editors never reached the standard set by recent BME doctoral graduation rates (40% female). In the aftermath of #metoo and considering new findings suggesting that higher proportions of female editors yields more female first- and last-authorships^13^, it appears that the field of BME is, as most other fields, is still in desperate need of policy revision to increase female representation on editorial boards.

On a similar note, we found that in BME, the percentage of Black journal editors (1%) was almost stagnant between 2016 and 2021. The need for spaces that are more supportive of Black^14^ and other racial minority groups is critical to improving retention, a statistic that is drastically reduced at graduate levels^15^. Additionally, advocating for greater proportions of Black people at elevated positions in science—like editorships—is also necessary to promote diversity at higher levels in science and BME^16^.

This study had two limitations. When calculating significance using χ^2^ analyses, we used an “expected” percentage of 40% when determining significant differences between female and male proportions. While 40% of doctoral BME degrees are awarded to women, the racial distribution of these awardees is not known. It is likely that these degrees are not equally awarded to women of White, Black, Asian, or Hispanic racial makeup; however, lack of data yielded our use of 40% as a standard across groups. Additionally, when calculating the ΔJIF, editors were deduplicated such that the ΔJIF with the highest magnitude was the journal each specific editor was associated with. This led to a loss of data that may have shown interesting trends in how many editors were associated with different journals that saw positive ΔJIF.

## Conclusion

The field of BME is widely regarded to be female- and URM-friendly, but analyses of gender and racial makeup at higher leadership levels show that BME may be more skewed towards White men than was previously considered. Although the proportion of women receiving undergraduate degrees in BME is higher than other fields of engineering, investigation into BME editorships show that important gatekeeping positions in the field are not as diverse, even in the aftermath of profound social movements like #metoo and BLM. Work to increase female and minority proportions at the editorial level must be prioritized to decrease biases in review and publication processes, and increase the diversity of BME throughout the pipeline.

## Acknowledgements

The authors appreciate the guidance of Dr. Fred Oswald, Rice University Department of Psychological Sciences, regarding the use of the R code to determine gender and racial probability.

## Notes

### Competing Interest Statement

The authors have declared no competing interest.

